# Microclimate, CO_2_ and CH_4_ concentration on Blue tits (*Cyanistes caeruleus*) nests: effects of brood size, nestling age and on ectoparasites

**DOI:** 10.1101/698340

**Authors:** Francisco Castaño-Vázquez, Santiago Merino, Soledad Cuezva, Sergio Sánchez

## Abstract

The presence of nestlings and other nest dwelling living beings in nests built in cavities could alter the composition of gases inside the cavity. In addition, this different concentration of gases could be used by some parasites as a cue to localize their hosts. Here, we explored the temporal variation in the concentration and isotopic signature of carbon dioxide (CO_2_) and methane (CH_4_) inside nest boxes of blue tits *Cyanistes caeruleus* during the nestling period (days 3, 8, 13, 20 and 21 post-hatching). Concentration of gases and isotopic signature were significantly different inside of nests than outside (forest) during the nestling period. CO_2_ concentration was higher inside nest while CH_4_ was lower than in forest air. The differences in the concentration of CO_2_ between nest boxes and forest were higher on days 8th and 20th of nestling age than in other ages while the CH_4_ was lower on day 20th of nestling age than in other ages. Moreover, CO_2_ variation was positive and significantly related with brood size and negative and significantly with hatching date. The difference of CO_2_ between inside of nests and forest on 8th day of nestling age were negative and significantly related to flea larvae abundance as measured at the end of the nestling period. A significant positive relationship was found between the difference of CH_4_ between nests and forest and the final abundance of flea larvae for the same nestling age. In addition, flea larvae abundance was positive and significantly related with the relative humidity in nests at 8 days of nestling age. Moreover, blowfly pupae abundance was negative and significantly related with the difference of temperature in nests at 3 day of nestling age. The condition of blue tit females was negative and significantly related with the abundance of blowfly pupae.

## Introduction

Although several effects of carbon dioxide (CO_2_) on organisms have been described long ago (see for example Leclerq et al. 2000, Allen et al. 2011), few studies have investigated how slight fluctuations in CO_2_ could affect them. For example, Bezemer and Knight (2001) demonstrated that the mixture of CO_2_ together with the increase of the temperature, led to a greater recruitment of juveniles and an advance in the emergence in common snail (*Helix aspersa*) during its development. Other organisms such as ants, bees and termites can use high CO_2_ concentration to locate their hives (Stange, 1996). It is also well known that mosquitoes, biting flies and ticks can use CO_2_ as a directional cue to find their hosts (Bogner, 1992, Takken and Knols, 1999) and that even slights fluctuations of CO_2_ can be used for that purpose (Grant et al. 1995, Mboera and Takken, 1997).

Several studies in birds have measured the concentration of CO_2_ during incubation period (Lamson and Edmond, 1914; Wangensteen et al. 1971; Walsberg, 1980) but few studies have investigated nest gas composition inside nests during nestling development. For example, White et al. (1978) found a higher concentration of CO_2_ in nests chambers of European bee-eater (*Merops apiaster*) than in outside air during nestling period and Mondain-Monval and Sharp (2018) found a higher concentration of CO_2_ in occupied burrows by Sand Martins (*Riparia riparia*) than in unoccupied burrows during nestling period. However, to our knowledge there are no works looking for the relationships between nest-gases (i.e. CO_2_ and CH_4_) and the abundance of parasites in nests. In addition, CH_4_ effects on birds and parasites in nests are completely unknown. The monitoring of methane (CH_4_) concentration and its isotopic signal can be used as good indicators of microbial activity in nests.

Geochemical tracing using the stable carbon isotopic signature δ^13^C is a very useful tool for understanding biogeochemical processes in ecosystems (Peterson and Fry, 1987). δ^13^C is the molecular ratio between the heavy carbon isotope ^13^C and the light one ^12^C (^12^C: 98,9 % and ^13^C: 1,1 %, Earth’s average). Every process, chemical reaction and phase change, involves a specific isotopic fractionation that determines the resulting ratio and confers a characteristic isotopic signal to the new generated phase. Thereby, the isotopic ratio δ^13^C introduces identifiers for the multiple carbon sources of production and mechanisms of transference in natural ecosystems (Cerling et al. 1991; Yakir and Sternberg, 2000; Maseyk et al. 2009; Garcia-Anton et al. 2017).

The application of δ^13^C isotopic ratio to avian ecology is a very powerful approach to infer diet and habitat selection (Inger and Bearhop, 2008). It has been widely used to characterizing populations and track migratory origins and dispersal patterns (Rubenstein and Hobson, 2004). Also, it has been frequently used to deduce composition of the diet in birds and reconstruct changes over time, including breath ^13^C/^12^C ratio analysis (Hatch et al. 2002). However, as far as we know, it has not yet been used previously for exhaustive characterization of bird habitats.

Microclimate inside cavity nest can vary considerably due to nest material composition, microbial activity, populations of arthropods, and birds’ activity. Nestling development adds different waste (i.e. feces, feathers debris) to nests, increase heat and changes in the relative humidity by evapotranspiration that could affect to the gas composition inside nests. All these changes can also affect attraction and development of parasites and other arthropods inside nests. However, our knowledge on how all these factors can interact among them to affect microclimatic conditions of nests and the life being developed in them is very scarce.

Here we study several factors potentially affecting composition of gases in nests and their relationship with the presence of nest-dwelling ectoparasite fauna. On the one hand, we study the variation of CO_2_ and CH_4_ and the stable isotope ratios of carbon (C) in both gases inside nests of blue tits *Cyanistes caeruleus* in relation to brood size, nestling age, temperature and relative humidity inside nests. On the other hand, we also test whether the concentration of gases inside nests differs from concentration on forest and whether this difference could be used by different ectoparasite species to detect their hosts thus showing greater abundances at the end of the nestling period.

## Material and methods

### Study population

This study was carried out during 2016 bird breeding season in a Pyrenean Oak, *Quercus pyrenaica* deciduous forest located in Valsaín (Segovia, central Spain, 40° 53’ 74 N, 4° 01’ W, 1200 m a.s.l). Blue tits *Cyanistes caeruleus* population breeding in wooden nest boxes has been studied since 1991 in this area (Sanz, 2002; Tomás et al. 2006). Nest boxes dimensions were of 17.5 cm height, 11.7 cm width, and 12.5 cm depth. The entrance hole was of 4.5 cm diameter and there are a little uncovered hollow (11.7 cm × 13 cm) just under roof for airing box. Nest material (plants, moss and feathers) occupied approximately 40% within nest box and nest space for air about 2559 cm^3^. Each breeding season, nest boxes are periodically inspected to determine some reproductive parameters like laying date, clutch size and hatching date (Merino, et al. 2000; Tomás, et al. 2007). This is an observational work that only implies ringing and measurement of birds and no other legal requirements are needed. Ringing permits to carry out this work were approved by Ignacio Quintanilla Rubio (Director General of the natural environment) and the Junta of Castilla y Leon (permit numbers: EP/SG/705/2015 and PNSG_SG_2015_0323). Our study area is part of the protected area of the National park of Sierra de Guadarrama (special protection area Montes de Valsaín).

### Measuring gases

Between May 24 and June 20, 2016, we carry out monitoring, sampling and study of 45 nest boxes occupied by blue tits. However, one desertion occurred during the study period for unknown reasons. As a result, the final sample size was 44 nest boxes. Firstly, the ecosystem was characterized from both climatic (meteorological data during the monitoring period of 2016: http://climaguadarrama.es/valsain/NOAA/NOAA-2016.txt) and geochemical points of view (CO_2_ and CH_4_ and δ ^13^C-CO_2_ y δ ^13^C-CH_4_ from oak forest air). For this purpose, we take samples of atmospheric air, soil air and nest-air during nesting period (from May 24 to June 20). A total of 192 nest-air samples were taken during nestling period at different nestling ages (44 samples at 3, 8, 13 and 21 posthatching and 16 samples of gases on day 20th of nestling age due the fact that nestlings had already fledged out from the other nests). Nest boxes were hanging from tree branches about 5 m above the ground. For gases collecting, nest boxes were carefully descended from tree branches and placed on ground. Next, gases extraction was carried out through the entrance hole of nest boxes and without opening them. After gas collection, nest boxes were open and number of nestling counted. Then next boxes were close again and located in the same position at the tree branch till the next sampling. Before the beginning of samplings from nest boxes occupied by birds, four nest boxes were sampled on May 18 at a similar time (7:00-9:00 AM), in order to have a reference value of the concentration of gases in nests built, but still without the presence of nestlings inside nests.

For the extraction of gases from nest boxes and soil, we use a micro-diagraph gas pump (KNF, NMP-830-KNDC - 12V, Neuberger, Freiburg; Germany) at 1.5 l·min^−1^ at atmospheric pressure. Gas extraction was carried out slowly and during the same time (20 sg) in order to obtain an amount of gas of approximately 0.5 liter from nest boxes. Similarly, atmospheric forest air was taken using a battery operated portable air pump (Aquanic S790, 0.010 Mpa 0.4 Lts/min Battery 1.5V; ICA, S.A, Spain). Gases samples were collected through a tube connected directly to the pump, while another tube of pump expelled the air and stored it in a plastic bag of 1 liter maximum capacity (Supelco; Bellefonte, PA 16823-0048 USA). These bags were sealed and subsequently analyzed in the laboratory before 24 hours. To analyze gas samples we used a spectrometer (Picarro G2201-i; 01-i; range: 100-4000 ppm Accuracy: + 0.05% 200 ppb measure (12C), 10 ppb + 0.05 % measured (13C), < 0.16 ‰ CO_2ati_. Patrick Henry Drive Santa Clara, California 95054; USA). This spectrometer uses technology CRDS (Cavity Ring-Down Spectroscopy) to identify and quantify the compounds contained in the analyzed air (Crosson 2008). Stable isotope ratios of carbon (C) are reported here in standard δ-notation with units of ‰ (δ ^13^C).

### Measuring temperature and relative humidity

Environmental conditions (i.e. temperature and relative humidity) were studied inside and outside of nests during nestling period (days 3, 8, 13 and 20 post-hatching). For this purpose, nest boxes were fitted inside with sensors that register both variables (iButton Hygrochron Temperature/Humidity Logger with 8KB Data-Log Memory DS 1923; 6 × 17 mm, temperature range: −20-85°C; resolution 0.0625°C; humidity range: 0-100% with a resolution 0.04%; Maxim IC, USA). Temperature and relative humidity were recorded every 45 min during nesting period. Sensors were located between the wall of the nest box and the rim of the nest. Similarly, four external sensors were located under empty nest boxes to register the temperature and relative humidity in the forest (HOBO U23-001; data logger; 10.2 × 3.8 cm, temperature range: −40-70°C; resolution 0.02°C; humidity range: 0-100% with a resolution 0.03%; Onset Data Loggers, Massachussetts, USA) in the study area during nesting period.

### Quantifying nest ectoparasites

Once nestlings fledged (day 21 post-hatching), sensors were removed and the nest material was collected in a sealed labelled plastic bag and transported to the laboratory to quantify ectoparasite abundance. Nests were stored at 4° C from 2 to 4 days and then were defaunated using Berlese funnels (see Tomas et al. 2007). Abundance of ectoparasites (mites and flea larvae) was estimated by examining the material obtained from funnels and counted under a magnifier glass microscope (OLYMPUS-SZX7; ACH1, Tokyo, Japan; see Merino and Potti, 1995). Nest material was then dismantled in search of blowfly larvae (*Protocalliphora azurea*). Moreover, the number of biting midges (*Culicoides* spp) and blackflies (Diptera: Simulidae) that parasitize these nests was estimated by using traps located inside the nest-boxes (see Tomás et al. 2008a). The traps consisting in a plastic petri dish (8.5 cm diameter; 55.67 cm^2^) containing a commercially available body gel-oil (Johnson’s baby chamomilla; Johnson & Johnson, Dusseldorf, Germany). The petri dishes were placed within nest boxes at day 10 post-hatching and retrieved at day 13. Afterwards, petri dishes were observed under magnifying glass to count the number of biting midges (*Culicoides* spp) and blackflies (Diptera: Simulidae) adhered to gel.

On the other hand, we also evaluate the effects of ectoparasites on the bird’s body condition (day 13 post-hatching). For this, seventy-five adults (37 males and 38 females) of blue tits were captured at 13 days post-hatching with traps mounted in nests-boxes. Adults were ringed with numbered aluminum rings when necessary. Nestlings were also measured and ringed at 13 days post-hatching. Mass of adults and nestling was measured with an electronic balance (± 0.1 g). Tarsus length of birds was measured with a digital caliper (± 0.1 mm) and wing length with a ruler (± 0.5 mm). Body condition index in adults and nestlings was calculated as the residuals of mass on tarsus length.

### Statistical analyses

Mann-Whitney’s test was used to test the differences between gases from nests and forest. In order to compare the differences of CO_2_ between inside nests and forest air at different nestling ages (days 3, 8, 13 and 20 post-hatching) we used General Linear Model (GLM). Differences of CO_2_ were transformed logarithmically in order to comply with normality assumptions. Then transformed differences of CO_2_ between nests and forest air were used as a dependent variable and nestling age as factor, including hatching date and brood size (number of nestlings) as covariates. Post-hoc test (Tukey HSD) was used in order to compare the differences of CO_2_ between different nestling ages. The same analysis and transformation were used with relative humidity.

Since the differences in the concentration of CH_4_ and temperature did not follow the normality assumptions nor even by transforming data, we used the test of Kruskal-Wallis in order to test for differences of CH_4_ and temperature between inside nests and forest air at different nestling ages (days 3, 8, 13, 20 post-hatching). For this analysis, we used as dependent variable the differences in the concentration of CH_4_ or temperature between nests and forest air and nestling age as factor. Wilcoxon’s test was used to test for paired differences in CH_4_ concentration or temperature between different nestling ages. Moreover, this same test was also used to test differences in CO_2_ and CH_4_ concentration between inside nests and forest air once nestlings had left the nest (day 21 post-hatching). Relationships between brood size and hatching date and differences in CH_4_ or temperature were tested separately by means of Spearman rank correlations.

To explore the relationships between gases and ectoparasites we used generalized linear models (GzLM) with a negative binomial distribution and log link function. Each ectoparasite abundance was used as dependent variable and the following independent variables were used: the differences in the concentration of CO_2_ and CH_4_ and the differences in temperature and relative humidity between inside of nests and forest, brood size, hatching date, and other ectoparasites (those not being used as dependent variable at each analysis). Then we use a likelihood-ratio chi-square test to compare the current model versus the null (in this case, intercept) model. A significant result indicates that model fitted to the data. In this case, we explore the significant independent variables.

In addition, we use multiple linear regressions to test for the relationships between different abundances of ectoparasites and body condition of adults and nestling blue tits. Graphics and statistical analyses were performed in STATISTICA 7 (<www.statsof.com>) and SPSS (IBM Corp. Released 2017. IBM SPSS Statistics for Windows, Version 25.0. Armonk, NY: IBM Corp).

### Data deposition

Data available from the Digital CSIC repository < >.

## Results

### Concentration of gases in the ecosystem

The geographical area of Valsain (Segovia) is characterized by a Mediterranean climate, a subtype of warm temperate climate with a dry and warm summer and cool mild winter (Csb climate type, Köppen-Geiger Classification slightly modified, AEMET-IM, 2011). Annual mean temperature in 2016 at the study area was 10.2°C and total annual precipitation was 556 mm (http://climaguadarrama.es/valsain/NOAA/NOAA-2016.txt). During the nesting period (from May 24 to June 19), there were 8 rainy days with total rainfall of 39.8 mm and the average temperature was 13.7°C (Table 1). Outside air CO_2_ mean values (forest air values) ranged from 411 to 547 ppm and the isotopic signal (δ^13^CO_2_) from −8.56 to −12.21‰. The local outdoor atmosphere has mean values of CO_2_ and CH_4_ concentrations slightly above the recent global monthly mean CO_2_ (roughly 405 ppm and 1.85 ppm, respectively, checked at https://www.esrl.noaa.gov/gmd/ccgg/data-products.html for our monitoring period). The highest CO_2_ values were recorded on the warmest days because of the increased flow from the soil during drier periods in which less soil humidity would induce an increase of the gas diffusivity (Werner et al. 2006, Garcia-Anton et al. 2012). CO_2_ mean values from soil ranged from 887 to 15025 ppm and the isotopic signature (δ^13^CO_2_) varies from −16.5 to −22.4‰. These data are consistent with the local climate and C3 vegetation. The CO_2_ / δ ^13^C-CO_2_ results of the samples analysis of forest air and soil air were studied using the Keeling approach. This method is widely used to characterize the δ1^3^C-CO2 of ecosystem respiration. The intercept values of the Keeling plot (δ^13^C_s_) for the samples collected over the monitoring period were −22.77‰. These values indicate a prevalence of C3 plant activity in this oak woodland (around −27‰, Amundson et al., 1998) plus a 4.4‰ diffusional enrichment from soil (Yakir and Sternberg, 2000).

**Table 1.** Average data of environmental conditions of the study area (Valsain; Segovia, central Spain, 40° 53’ 74 N, 4° 01’ W, 1200 m a.s.l) during nesting period.

| Date | T (°C) | Rain (mm) | Forest |  |  |  | Soil |  |  |  |
| --- | --- | --- | --- | --- | --- | --- | --- | --- | --- | --- |
| | | | $\text{CO}_2$ (ppm) | $\delta^{13}\text{CO}_2$(‰) | $\text{CH}_4$ (ppm) | $\delta^{13}\text{CH}_4$(‰) | $\text{CO}_2$ (ppm) | $\delta^{13}\text{CO}_2$(‰) | $\text{CH}_4$(ppm) | $\delta^{13}\text{CH}_4$(‰) |
| 24-may-16 | 14.9 | 0 | 411 | -9.39 | 1.97 | -51.55 | 15025 | -21.39 | 0.20 | -26.52 |
| 25-may-16 | 12.1 | 0 | 442 | -9.35 | 2.00 | -54.91 | 3194 | -21.13 | 1.39 | -50.56 |
| 26-may-16 | 13.6 | 0 | 480 | -10.61 | 2.07 | -54.09 | 9671 | -22.26 | 0.13 | -51.87 |
| 27-may-16 | 12.9 | 0 | 428 | -8.56 | 2.09 | -61.27 | 8846 | -21.99 | 0.14 | - |
| 28-may-16 | 9.5 | 6 | 428 | -9.28 | 1.98 | -51.50 | 7919 | -21.94 | 0.32 | -12.84 |
| 29-may-16 | 8.7 | 10.2 | 445 | -9.21 | 2.12 | -62.94 | 9253 | -22.34 | 0.12 | - |
| 30-may-16 | 9.3 | 1.2 | 424 | -9.25 | 2.02 | -53.50 | 3108 | -21.54 | 1.44 | -46.30 |
| 31-may-16 | 10.3 | 0 | 439 | -10.05 | 2.13 | -54.23 | 10845 | -22.42 | 0.13 | -10.82 |
| 01-jun-16 | 11.5 | 0 | 424 | -9.63 | 2.00 | -50.71 | 9910 | -22.25 | 0.17 | -31.32 |
| 02-jun-16 | 13.2 | 0 | 473 | -11.06 | 1.98 | -53.51 | 9523 | -22.16 | 0.20 | -15.88 |
| 03-jun-16 | 15.5 | 0 | 488 | -10.31 | 2.06 | -57.98 | 8686 | -21.93 | 0.19 | -48.26 |
| 04-jun-16 | 14.5 | 0 | 436 | -9.64 | 2.09 | -55.45 | 8257 | -21.84 | 0.20 | -17.13 |
| 05-jun-16 | 15.5 | 0 | 447 | -9.93 | 2.00 | -53.42 | 8004 | -21.93 | 0.18 | - |
| 06-jun-16 | 16 | 0 | 487 | -11.34 | 2.02 | -56.18 | 7616 | -21.88 | 0.24 | -26.96 |
| 07-jun-16 | 17.8 | 0 | 463 | -10.32 | 2.05 | -56.12 | 4003 | -19.21 | 1.04 | -32.12 |
| 08-jun-16 | 19.4 | 0 | 542 | -12.21 | 2.03 | -55.03 | 3056 | -20.90 | 1.24 | -42.99 |
| 09-jun-16 | 19.7 | 0 | 547 | -12.08 | 2.00 | -54.65 | 3819 | -21.28 | 1.02 | -42.89 |
| 10-jun-16 | 16.7 | 0.2 | 456 | -10.92 | 2.04 | -56.06 | 1907 | -19.39 | 1.45 | -45.41 |
| 11-jun-16 | 16 | 0 | 460 | -10.6 | 2.01 | -55.09 | 887 | -16.51 | 1.75 | -45.11 |
| 12-jun-16 | 16.4 | 0 | 474 | -10.77 | 2.04 | -55.58 | 1518 | -18.90 | 1.56 | -45.32 |
| 13-jun-16 | 18.1 | 0 | 448 | -10.50 | 1.99 | -55.31 | 1666 | -19.19 | 1.52 | -45.33 |
| 14-jun-16 | 14.9 | 0 | 424 | -10.22 | 1.97 | -50.64 | 4790 | -21.43 | 0.63 | -34.64 |
| 15-jun-16 | 10.1 | 9.4 | 430 | -10.09 | 1.95 | -53.44 | 3565 | -21.08 | 0.96 | -47.87 |
| 16-jun-16 | 7.9 | 11.4 | 429 | -10.27 | 1.94 | -53.10 | 3319 | -21.47 | 1.16 | -47.94 |
| 17-jun-16 | 10.3 | 1.2 | 435 | -9.99 | 1.99 | -56.64 | 3085 | -21.19 | 1.32 | -52.11 |
| 18-jun-16 | 11.3 | 0.2 | 427 | -9.91 | 2.03 | -53.34 | 8155 | -21.88 | 0.27 | -36.09 |
| 19-jun-16 | 11.9 | 0 | 437 | -10.11 | 2.01 | -54.95 | 8375 | -21.52 | 0.18 | -33.06 |
| 20-jun-16 | 16.3 | 0 | 411 | -8.97 | 1.96 | -58.19 | 8259 | -21.51 | 0.18 | -40.24 |

Temperature (°C), CO<sub>2</sub> concentration (ppm), CO<sub>2</sub> isotopic signal ( $\delta^{13}\text{C}_{\text{CO}_2(\text{‰})}$ ), CH<sub>4</sub> concentration (ppm), and CH<sub>4</sub> isotopic signal ( $\delta^{13}\text{C}_{\text{CH}_4(\text{‰})}$ ) are shown. Data from June 20 correspond to nests without nestlings.

### Concentration of gases, age, brood size and hatching date

At the start of our sampling (May 18, 2016), we did not observe significant differences between the concentration of CO_2_ inside nest boxes and forest air that containing nests although without nestlings (unpaired *t*-test: *t* = 1.27, df = 5, p = 0.259; Table 2). However, there are significant differences for isotopic signal (δ^13^CO_2(‰)_)(unpaired *t*-test: *t* = −2.90, df = 5, p = 0.033; Table 2). CH_4_ concentration was significantly lower inside those nests as compared to the forest air (unpaired *t*-test: *t* = −5.73, df = 5, p = 0.002). However, there are no significant differences for the isotopic signal (δ^13^CH_4(‰)_; unpaired *t*-test: *t* = 2.27, df = 5, p = 0.072). Likewise, all nest samples with nestlings during nesting period (from 24 May to June 20) shown higher concentration of CO_2_ and a lighter isotopic signal (δ^13^CO_2 (‰)_) than forest air (Mann-Whitney test for CO_2_: *Z* = −8.11, p < 0.001; Mann-Whitney test for δ^13^CO_2(‰)_: *Z* = 8.22, p < 0.001, Table 3). However, we observed a lower concentration of CH_4_ and a heavier isotopic signal (δ^13^CH_4(‰)_) in nests with nestlings than in forest air (Mann-Whitney test for CH_4_: *Z* = 7.90, p < 0.001; Mann-Whitney test for δ^13^CH_4 (‰)_: *Z* = 7.79, p < 0.001, Table 3; Fig. 1).

**Figure 1.**
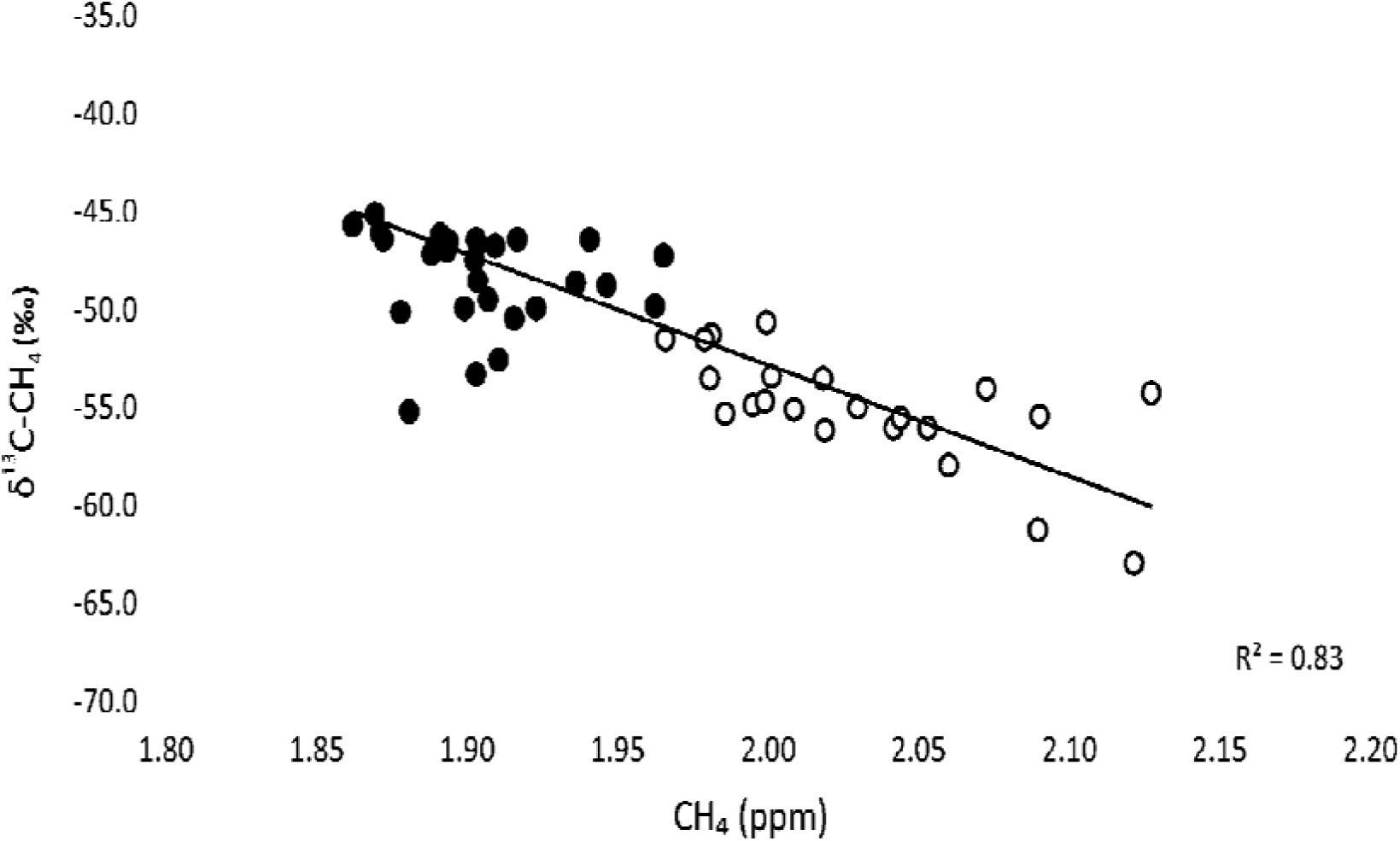
Concentration values of methane (CH_4_) and δ13C-CH4 from both the samples taken at the nests and from the outdoor air samples throughout the monitoring period.

**Table 2.** Average values of concentration (ppm) and isotopic signals (‰) of CO_2_ and CH_4_ from nestboxes and forest.

| Date | Nestbox |  |  |  | Forest |  |  |  |
| --- | --- | --- | --- | --- | --- | --- | --- | --- |
| | CO<sub>2</sub> | $\delta^{13}\text{C-CO}_2$ | CH<sub>4</sub> | $\delta^{13}\text{C-CH}_4$ | CO<sub>2</sub> | $\delta^{13}\text{C-CO}_2$ | CH<sub>4</sub> | $\delta^{13}\text{C-CH}_4$ |
| 18 May | 410 | -8.82 | 1.95 | -48.70 | 405 | -8.34 | 1.98 | -51.28 |
| 10 June | 441,74 | -10,90 | 1,87 | -45,76 | 455,89 | -10,92 | 2,04 | -56,06 |
| 11 June | 429,61 | -10,19 | 1,88 | -46,39 | 460,20 | -10,63 | 2,01 | -55,09 |
| 13 June | 420,81 | -10,00 | 1,88 | -46,44 | 447,74 | -10,50 | 1,99 | -55,31 |
| 14 June | 431,48 | -10,91 | 1,89 | -54,36 | 423,67 | -10,22 | 1,97 | -50,64 |
| 15 June | 439,27 | -10,89 | 1,88 | -50,18 | 429,97 | -10,09 | 1,95 | -53,44 |
| 16 June | 458,72 | -12,08 | 1,90 | -52,18 | 429,13 | -10,27 | 1,94 | -53,10 |
| 17 June | 461,18 | -11,17 | 1,91 | -50,89 | 434,54 | -9,99 | 1,99 | -56,64 |
| 18 June | 466,10 | -11,23 | 1,91 | -53,41 | 427,23 | -9,91 | 2,03 | -53,34 |
| 19 June | 433,78 | -10,65 | 1,91 | -52,57 | 436,99 | -10,11 | 2,01 | -54,95 |

Data from nestboxes with nests but without nestlings and from samples of forest air taken at the same dates. Data from the 18 of May are from nests previous to the breeding season and data from June are from nests once nestlings fledge.

**Table 3.** Average values and standard deviation (SD) of the concentration (ppm) and isotopic signal (‰) of CO_2_ and CH_4_.

| Gases | Nest boxes (n=148) |  | Forest (n=27) |  | Mann-Whitney test |  |
| --- | --- | --- | --- | --- | --- | --- |
|  | mean | SD | mean | SD | □ | p |
| CO<sub>2</sub> (ppm) | 1065.06 | 756.21 | 452.74 | 33.68 | -8.11 | < 0.001 |
| CH<sub>4</sub> (ppm) | 1.91 | 0.03 | 2.02 | 0.04 | 7.90 | < 0.001 |
| $\delta^{13}\text{CO}_2$ (‰) | -17.32 | 3.21 | -10.20 | 0.84 | 8.22 | < 0.001 |
| $\delta^{13}\text{CH}_4$ (‰) | -47.49 | 2.39 | -54.84 | 2.74 | 7.79 | < 0.001 |

Data measured in nest boxes of blue tits (*Cyanistes caeruleus*) at different nestling ages and from outside of nest (forest) from May 24 to June 19, 2016.

Once nestlings fledge, differences in concentration of CO_2_ were not significant (U-Mann Whitney: Z= 0.75, p = 0.45 comparing average data from nests and forest measured each day, N= 9). However, there are significant differences for isotopic signal (δ^13^CO_2(‰)_)(U-Mann Whitney: Z= −2.16, p = 0.03). CH_4_ concentration was significantly lower and the isotopic signal (δ^13^CH_4 (‰)_) significantly heavier inside nests once nestlings fledge as compared to the forest air (paired *t*-test: *t* = −7.97, df = 8, p < 0.001 and paired *t*-test: *t* = 2.58, df = 8, p = 0.03 respectively; see Table 2).

For simplification here we present only results concerning the difference in concentration of gases between nests and forest air in relation with other variables of study (brood size, hatching date, temperature, relative humidity and ectoparasites). Results by using other variables are similar to those presented here. We decide to select differences in gases concentration between nest boxes and forest because it could be the cue used by some ectoparasites to localize nestlings. The lack of clear differences between forest and nest boxes gas concentrations will render a lack of signal for parasites. Differences in the concentration of CO_2_ between nest boxes and forest air differ with nestling age (ANCOVA, F_3, 141_ = 19.32, p < 0.001 Fig. 2a) with a maximum difference by the age of 20 days when nestlings begin to leave the nest. Moreover, the difference of concentration of CO_2_ was positive and significantly related with the brood size (ANCOVA, F_1, 141_ = 8.25, p = 0.004) and negative and significantly with the hatching date (ANCOVA, F_1, 141_ = 6.32, p =0.013). Post hoc analyses show that differences in the concentration of CO_2_ between nests and forest air differ significantly between all ages except for differences between days 3 and 13 of nestlings’ age (Tukey test: p < 0.05 for all cases). On the other hand, we did not find significant differences between the concentration of CO_2_ inside nests and forest air once nestlings had left the nest (day 21 post-hatching; Wilcoxon test: n = 44, Z = 1.55, p = 0.120).

**Figure 2a.**
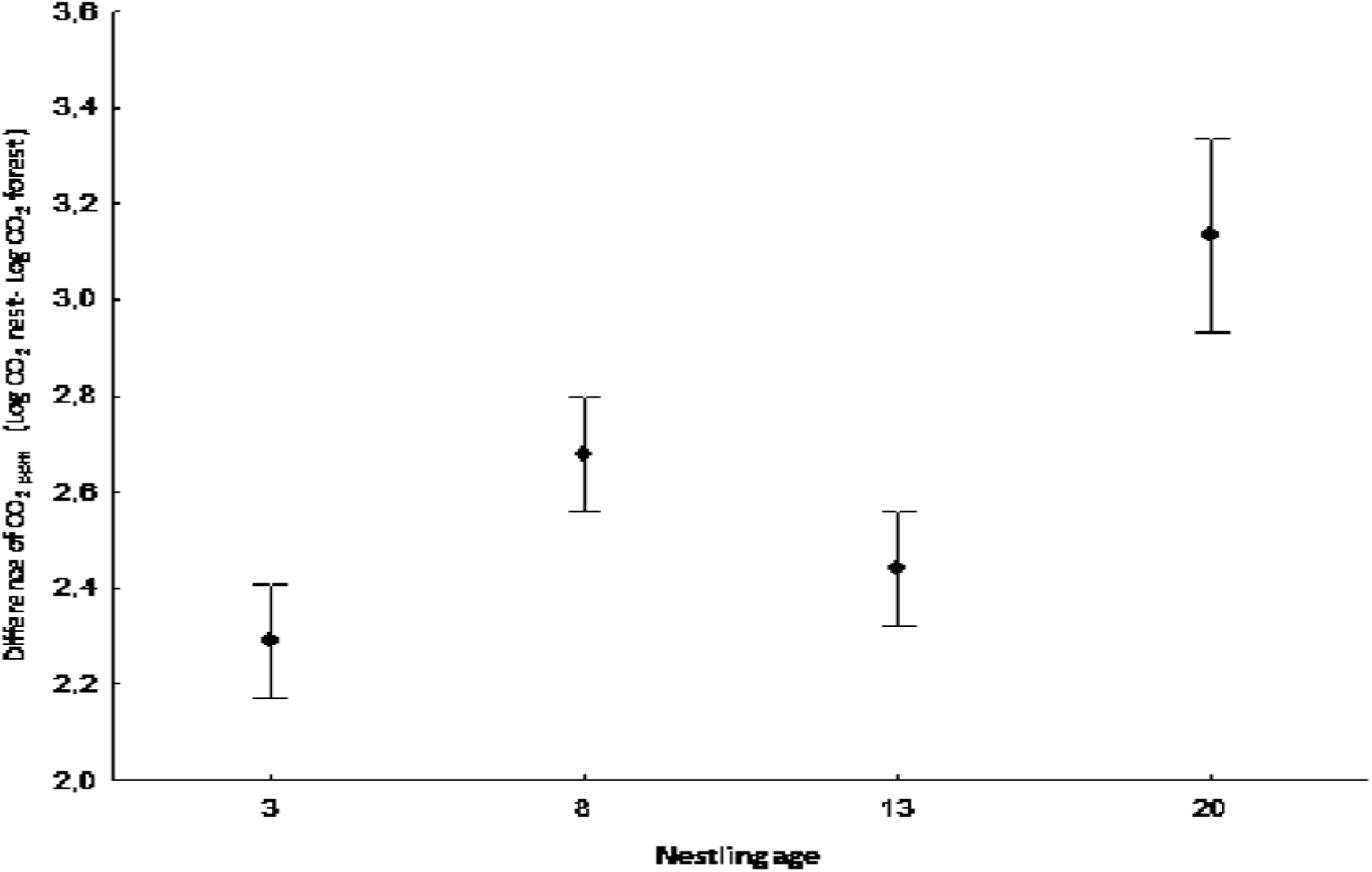
ANCOVA to differences of concentration in CO_2_ between nests and forest air at different nestling ages (days 3, 8, 13 and 20). Concentrations in ppm. Bars show mean ± SE for each gas.

The difference in CH_4_ concentration between nest boxes and forest air showed significant differences with nestling age (Kruskal-Wallis test: χ^2^ = 13.05, df = 4, p < 0.005 Fig. 2b) but this time with a minimum for day 20. However post-hoc test only show significant differences between days 13 and 20 of nestlings’ age (p < 0.005). Moreover, we did not find a significant relationship between the difference of concentration of CH_4_ and the brood size (Spearman, n = 145, *r_s_* = −0.11, p = 0.182) or hatching date (Spearman, n = 145, *r_s_* = −0.09, p = 0.254). Likewise, we found a significant lower concentration of CH_4_ inside nests than in forest air once nestlings had left the nest (day 21 post-hatching; Wilcoxon test: n = 44, Z = 5.77, p < 0.001).

**Figure 2b.**
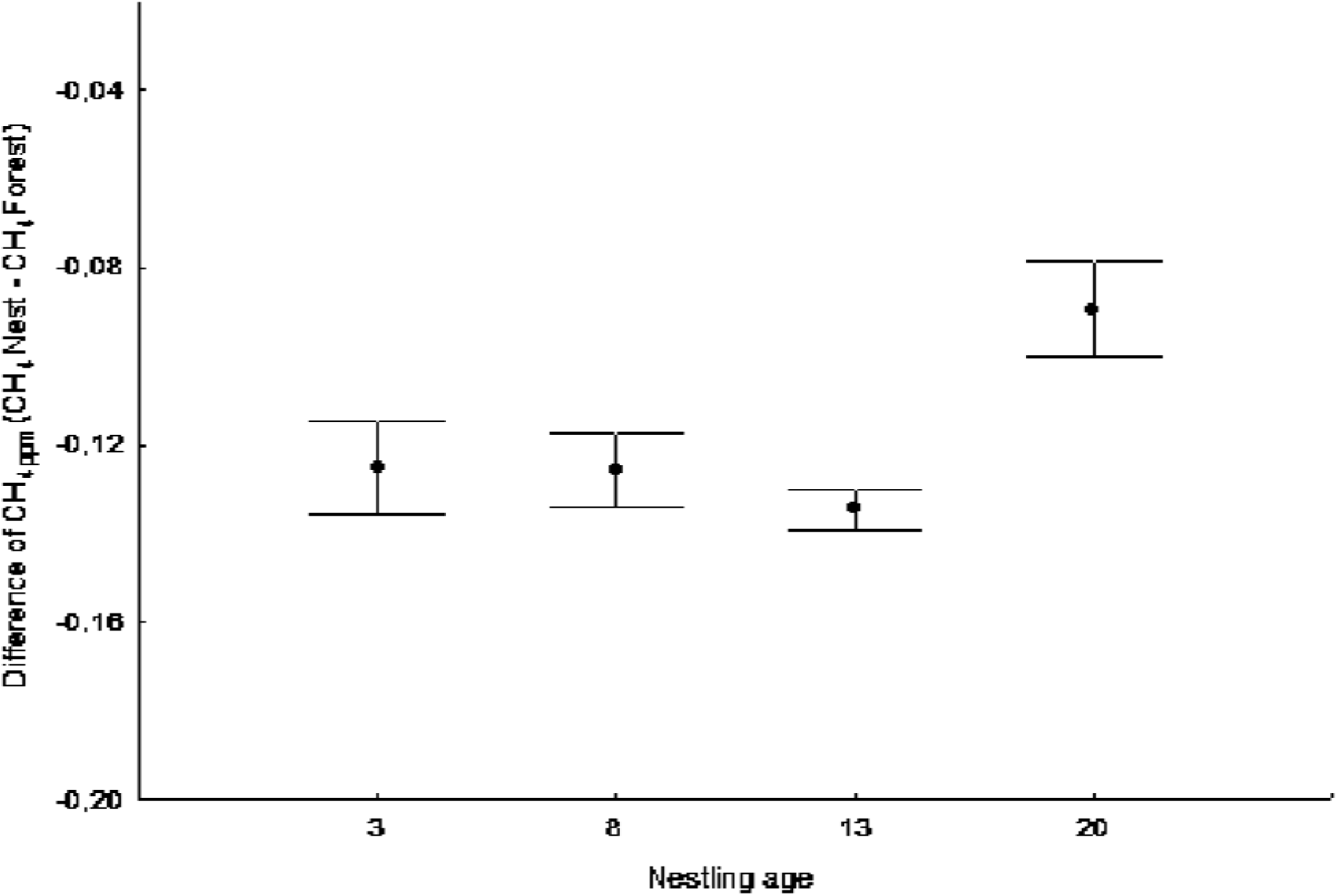
Kruskal-Wallis ANOVA to differences of concentration in CH_4_ between nests and forest air at different nestling ages (days 3, 8, 13 and 20). Concentrations in ppm. Bars show mean ± SE for each gas.

The difference in CH_4_ concentration between nest boxes and forest air showed significant differences with nestling age (Kruskal-Wallis test: χ^2^ = 13.05, df = 4, p < 0.005 Fig. 2b) but this time with a minimum for day 20. However post-hoc test only show significant differences between days 13 and 20 of nestlings’ age (p < 0.005). Moreover, we did not find a significant relationship between the difference of concentration of CH_4_ and the brood size (Spearman, n = 145, *r_s_* = −0.11, p = 0.182) or hatching date (Spearman, n = 145, *r_s_* = −0.09, p = 0.254). Likewise, we found a significant lower concentration of CH_4_ inside nests than in forest air once nestlings had left the nest (day 21 post-hatching; Wilcoxon test: n = 44, Z = 5.77, p < 0.001).

### Temperature, relative humidity and concentration of gases

The difference of temperature between inside nest boxes and forest air differ with nestling age (Kruskal-Wallis test: χ^2^ = 25.25, df = 3, p < 0.001, Fig. 3a) with a maximum difference by the age of 13 days. Post-hoc test show that significant differences exist between different ages except for days 8 and 13 and days 3 and 20 of nestling age (p > 0.05 for both cases). Forest air temperature increased from 3rd day of nestling age reaching a maximum at 13 days of nestling age (see Fig. 3a). Likewise, the difference of temperature between nest and forest air was not significantly related with brood size (Spearman, n = 145, *r_s_* = −0.13, p < 0.119). However, hatching date was related negative and significantly with the difference of temperature between inside nest and forest air with later hatched nest showing lower difference in temperature (Spearman, n = 145, *r_s_* = −0.30, p < 0.005).

**Figure 3a.**
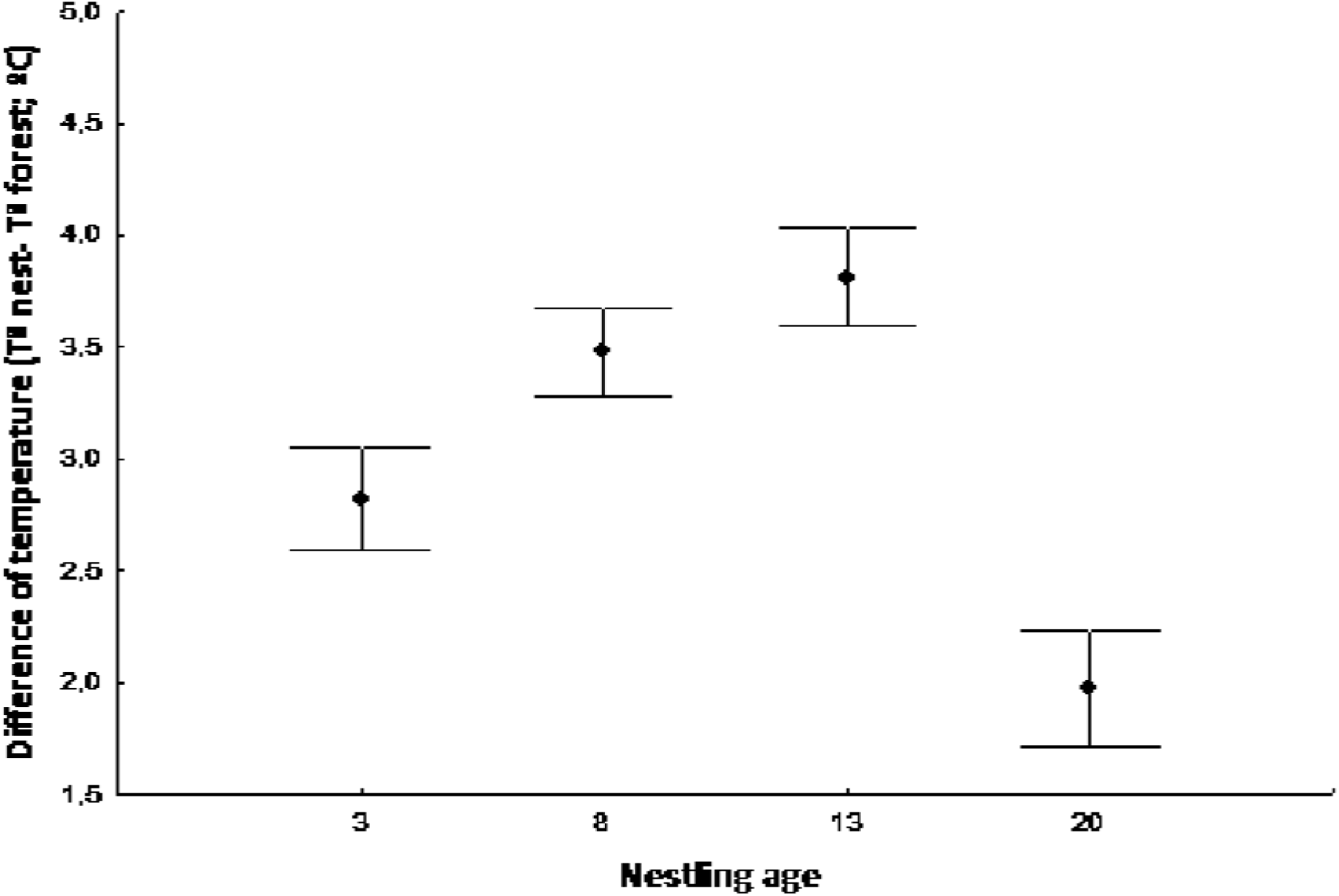
Average difference of temperature between nests and forest air at different nestling ages. Bars show mean ± SE for each gas.

The difference of relative humidity between nest boxes and forest air differ with nestling age (ANCOVA, F_3, 139_ = 10.12, p < 0.001 Fig. 3b) with a minimum difference at 13 days of nestling age, when temperature differences between nest boxes and forest air were higher (Fig. 3a). Moreover, the difference of relative humidity was positive and significantly related with the brood size (ANCOVA, F_1, 139_ = 4.26, p = 0.040) and with the hatching date (ANCOVA, F_1, 139_ = 4.89, p =0.028). Post-hoc tests show significant differences for the difference of relative humidity between ages 3 and 8; 3 and 13; 3 and 20 (Tukey test: p < 0.05 for all cases).

**Figure 3b.**
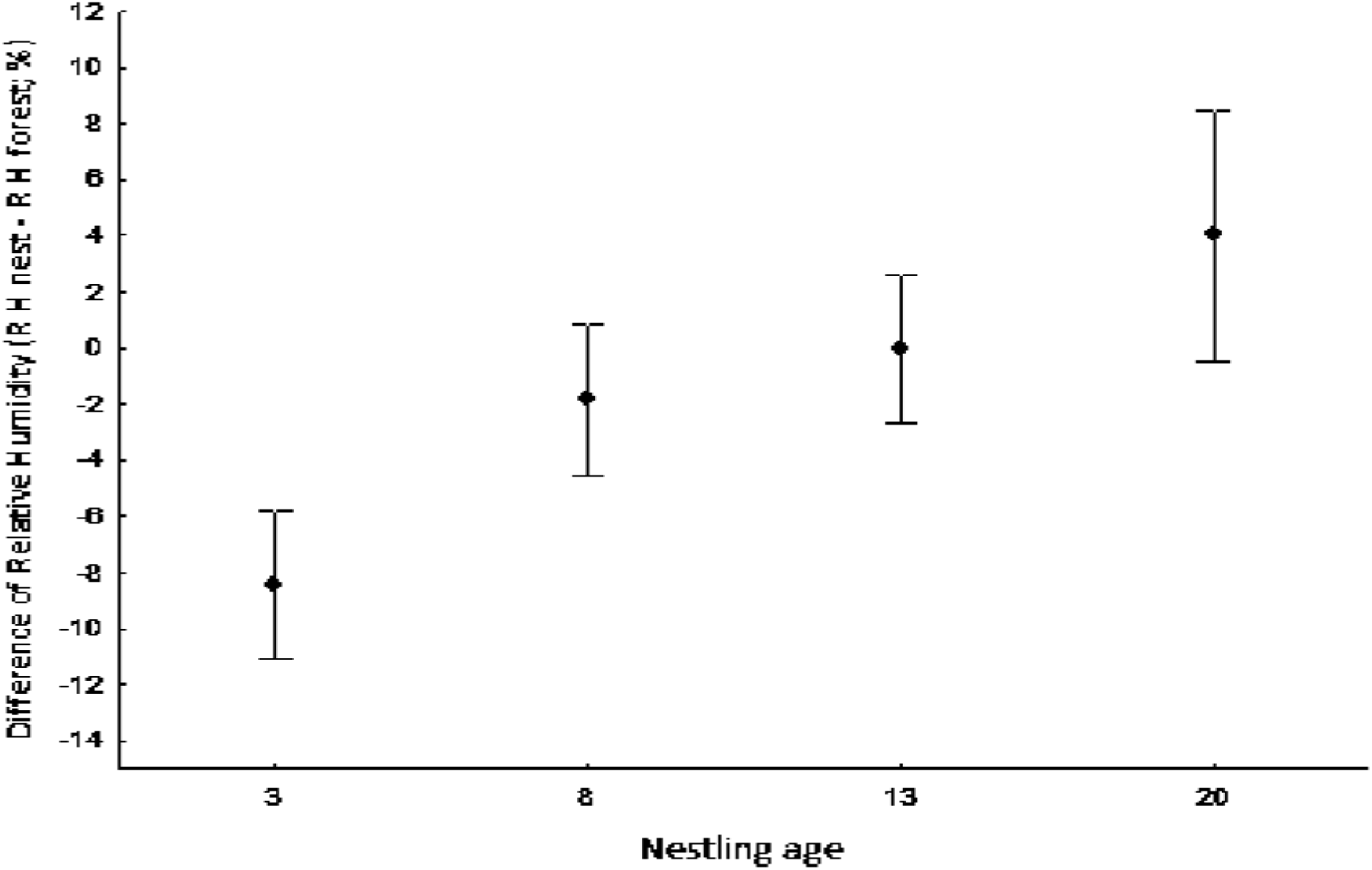
Average difference of relative humidity between nests and forest air at different nestling ages. Bars show mean ± SE for each gas.

On the other hand, the differences in the concentration of gases (CO_2_ and CH_4_) between nests and forest showed different relationships with the differences of temperature and relative humidity at different nestling age. The differences in the concentration of CH_4_ between nest boxes and forest air was negative and significantly related with the difference of relative humidity between nests and forest air at 3 and 20 days of nestling age (Spearman, p < 0.05, for both cases). Conversely, this same relationship was positive and significant at 8 days of nestling age (Spearman, n = 42, *r_s_* = 0.42, p < 0.01). Moreover, the differences in concentration of CH_4_ between nest boxes and forest air was positive and significantly related with the differences of temperature at 3 days of nestling age (Spearman, n = 41, *r_s_* = 0.31, p = 0.045). Likewise, we also found a significant and negative relationship between the difference of concentration of CO_2_ between nest boxes and forest air with the relative humidity at 13 days of nestling age (Spearman, n = 43, *r_s_* = −0.31, p = 0.038).

### Nest ectoparasites

The difference in the concentration of CO_2_ between nests and forest on 8th day of nestling age was negative and significantly related to the abundance of flea larvae (n= 41, F_1, 30_ = 20.6, p < 0.001). However, the difference in the concentration of CH_4_ between nests and forest the same day was positive and significantly related to the abundance of flea larvae (n = 41, F_1, 30_ = 4.9, p = 0.034). Likewise, there are several relationships between different ectoparasites when the effect of gases at different ages were taking into account: blowfly pupae abundance was negative and significantly related with the abundance of mites at 3 day of nestling age (n= 40, F_1, 29_ = 7.1, p = 0.012); Final abundance of biting midges was positive and significantly related with the abundance final of blackflies at 3, 8 and 13 days of nestling age (p < 0.01, for all cases); the abundance of flea larvae was negative and significantly related with the abundance of blackflies inside nests at 8 days of nestling age (n= 41, F_1, 30_ = 9.8, p = 0.004) and the same day of age, the abundance of flea larvae was positive and significantly related with the abundance of mites and blowfly pupae inside nests (n= 41, F_1, 30_ = 6.4, p = 0.016; n= 41, F_1, 30_ = 6.8, p = 0.014 respectively).

The difference of temperature between nests and forest on 3th day of nestling age was negative and significantly related to the abundance of blowfly pupae (n= 40, F_1, 29_ = 10.9, p = 0.003). On 8th day of nestling age, the abundance of flea larvae was positive and significantly related with the relative humidity (n= 41, F_1, 30_ = 4.2, p = 0.048). The same day, the abundance of flea larvae was negative and significantly related with the brood size (n= 41, F_1, 30_ = 14.9, p = 0.001). On 13th day of nestling age, the abundance of biting midges was positive and significantly related with the hatching date (n= 42, F_1, 31_ = 5.2, p = 0.028).

Only blowfly pupae were related negatively and significantly with female body condition (Multiple linear regression, F_1, 33_ = 7.0, p = 0.012). Condition of males or nestlings did not relate significantly with the abundance of any ectoparasite (p > 0.05; in all cases).

## Discussion

The concentration of gases (i.e. CO_2_ and CH_4_) inside the breeding cavities of birds is poorly studied and much less the effects of these gases on the attraction to and growing of parasite populations in the nests. In this study, we showed that CO_2_ and CH_4_ concentration and isotopic signal (δ^13^CO_2 (‰)_ δ^13^CH_4 (‰)_) inside nest boxes of blue tits *Cyanistes caeruleus* were different from forest air during nesting period (from 24 May to June 20) in an ecosystem dominated by the activity of C3 plants. Nest-air shown a higher concentration of CO_2_ and lighter δ^13^CO_2 (‰)_ than forest air. In the same way, White et al. (1978) found a higher concentration of CO_2_ in nests chambers of European bee-eater (*Merops apiaster*) than outside air during nestling period. In addition, Mondain-Monval and Sharp (2018) found a lower concentration of CO_2_ in unoccupied burrows of Sand Martins (*Riparia riparia*) than in occupied burrows. Before and after nestling period, we did not find significant differences between the concentration of CO_2_ in nests without nestlings and forest air. Thus, as expected, the presence of nestlings increases the concentration of CO_2_ inside nests. Slight but significant differences in concentration of gases in absence of nestlings clearly shown that nesting material include microbial activity inside nests.

In our study, the higher differences in CO_2_ concentration between nest-air and forest air were observed at 8 and 20 days of nestling age. It is possible that differences in the concentration of CO_2_ observed on 8th day of age were due to increase of growth rate (>mass) after hatching. This fact could have increased the metabolic rate of nestlings and therefore a higher accumulation of CO_2_ inside nests. In this sense, Morganti et al. (2017) showed that blue tit nestlings have a higher mass gain between 4th and 8th day of age. In addition, it is possible that nestlings increased their metabolic rate due to the beginning of thermoregulatory development, a process that for passerine birds usually occurs around 8 days of age (Visser, 1998, Pereyra and Morton, 2001). Moreover, the highest differences in the concentration of CO_2_ between nests and forest were found on 20th day of nestling age. It is possible that highest differences during this day were due to the movement of nestlings inside nests during exercise of wings just before leaving the nest. In the same way, Wickler and Marsh (1981) found a positive relationship between CO_2_ and nestling age in nests chambers of bank swallow (*Riparia riparia*). In addition, Mersten-Katz et al. (2012) found that the rate of CO_2_ production had little influence on the gas composition inside nests during early stages of nestling development, being this rate higher about 15 days of nestling age due to activity of nestlings within the nests. In our study, we also found that the difference in CO_2_ concentration between nests and forest air was positive and significantly related with the brood size. Obviously, it is expected that more nestlings would produce more CO_2_. In addition, later hatched nestlings produce lower differences in concentrations of CO_2_. This may be due to different factors including more or less complex effects of other climatic variables. For example, difference in temperature inside nests was lower for later hatched nestlings and lower relative humidity inside nests was related with higher difference in CO_2_ when nestlings had 13 days of age. On the other side, Howe et al. (1987) suggested that due to the presence of nestlings of Northern flickers (*Colaptes auratus*) inside nests, there was a increase of heat that led to a higher accumulation of CO_2_. However, we fail to find a significant relationship between differences in CO_2_ and differences in temperature in blue tit nests.

However, the concentration of CH_4_ was lower and the isotopic signal (δ^13^CH_4 (‰)_) significantly heavier inside nest boxes than in forest air and this difference is clear even in nest without nestlings. This must be indicative of the activity of methanotrophic oxidizing bacteria (MOB) inside nest boxes from the very beginning of birds breeding activity. Differences in concentration of CH_4_ show low variation with nestling age except when they reach 20 days of age. It is possible that this lower difference of CH_4_ was due to an increase in activity of MOB living in nests. MOB activity seems be the main responsible for consumption of CH_4_ (Hanson and Hanson, 1996) and in the nests this process could be intensified and slightly modified as a result of the insulating effect on the boxes and the metabolic activity of the birds inside (probably related to accumulation of waste and faeces). However, we do not find effects of brood size or hatching date on CH_4_ differences thus indicating that methanotrophic activity inside nest may be linked to nest materials more than to nestling activity. Lower concentration of CH_4_ and higher isotopic signal for this gas in nests imply that oxidization of the gas is occurring in nests reducing its concentration. In addition, the isotopic signal changes to heavier values thus indicating that methanotrophic oxidizing bacteria (MOB) are responsible of the oxidization process (see Figure 1). This process is intensified and slightly modified as a result of the insulating effect of the boxes and the metabolic activity of the birds inside (probably related to accumulation of waste and feces). In this respect we find some variation in difference of CH_4_ with temperature and relative humidity but the pattern is not always clear. For example, we find increases and decreases of differences in CH_4_ concentration in relation to relative humidity depending of the age of nestlings and the positive relationship with temperature only appear when nestlings have 3 days of age. This may be indicative of the need of a minimum temperature for optimal bacterial growing but in any case once again differences in methane do not appear clearly related with variables related with nestlings like age or brood size. Finally, soil CH_4_ concentration was always lower and its isotopic ratio was always lighter than values obtained in nests and exterior air (see Table 1). These data are consistent with usual removal of methane by bacterial oxidation in aerobic soils (Conrad 1996) and the subsequent consumption of the atmospheric methane in the ecosystem.

We do not know of studies that have investigated temperature variation inside nests at different nestling ages. Temperature difference between nest boxes and forest air was higher when nestlings had 13 days of age. Similarly, the difference of relative humidity inside nest boxes and forest air was lower, almost zero, the same day of age. A higher difference of temperature during this day could have been caused by a warmer external environment (forest air). In fact, we found that forest air temperature increased from 3rd day of nestling age to a maximum at 13 days of age. These results matched with the reduction in relative humidity inside blue tits nests after experimental modification of nest temperature found by Castaño-Vázquez et al. (2018). In addition, we find significant positive relationships between differences in relative humidity and brood size and hatching date. The later could be related with the rainy days at the end of the breeding season. On the other hand the presence of a higher number of nestlings in nests could also increase transpiration and produce a higher relative humidity inside nest.

One of our main predictions was that differences in gases concentrations between inside nests and forest could be used as a cue for host detection by ectoparasites. The significant difference in concentration of both CO_2_ and CH_4_ between nests and forest support that possibility although differences in CH_4_ appear to be less clearly related with the presence of nestlings in nests. However, our results do not show clearly that parasites used differences in gas concentration to locate their blue tit hosts. We only find significant relationships between differences in concentration of gases and abundances of fleas in nests, a negative one with CO_2_ and a positive one with CH_4_. However, fleas use to reach nests by attaching to adult birds when they visit old nests or cavities containing fleas (Marshall 1981). Thus the relationship of fleas with gases may be related to a direct effect of gases on flea development or survival. In this respect Downs et al. (2015), observed a reduction in the activity and survival of fleas when exposed to a higher level of CO_2_. The positive relationship between flea abundance and differences in CH_4_ concentration is difficult to explain although could be due to the use of nest debris as food for both flea larvae and methanotrophic bacteria. The lack of relationships between differences in gas concentration and abundances of other parasites could be due to the implication of other factors in host detection by these parasites. For example, Bishop et al. (2008) argued that initial attraction of biting midges to host was first visual, and host odours (i.e. CO_2_ and 1-octen-3-ol) were secondary and occurred once host was located. Alternatively, Tomás and Soler (2016) also proposed that acoustic signals produced during nestling begging behaviour could attract blood-feeding insects to nests.

We also find positive and negative relationships between different ectoparasite abundances when looking for relationships between ectoparasites and gases. For example blood feeding flying insects are positively related indicating that blackflies and biting midges could use similar cues to find their hosts or that some nests are more susceptible to parasitism than others (e.g. by being closer to insect oviposition sites or due to differences in antiparasite behaviours of birds; Tomás et al. 2008a). Fleas also appear associated positively with mite and blowfly abundances but negatively with blackflies. The positive associations in this case could be related with the use of the same food source, nestlings, but the negative relationship with blackflies is difficult to explain. However, some studies have shown that arthropod abundance inside nests can affect negatively flea abundance (Heeb et al. 2000, Hanmer et al. 2017). We also find a negative relationship between final abundance of blowflies and the final abundance of mites in nests. This relationship was previously reported by Merino and Potti (1996) in pied flycatchers *Ficedula hypoleuca* and explained as the result of competition between both parasites for resources. Other studies have shown that the presence of arthropods in nests can reduce the abundance of mites (Dube et al. 2018).

Temperature and humidity are important factors for ectoparasite development. For example several studies have shown that an environment with higher humidity could favour the increase of the number ectoparasites (Heeb et al. 2000; Moyer et al. 2002). Thus the positive relationship between relative humidity and flea larvae abundance inside nests could be explained by the importance of humidity for them. We also found a negative relationship between difference in temperature and the abundance of blowfly pupae. In this respect, an experimental increase of temperature inside nest produce a reduction in the number of blowfly pupae in blue tit nests (see Castaño-Vázquez et al. 2018). In addition, Mennerat et al. (2019) found a lower abundance of blowflies in blue tit nests due to effect of warmer summer during previous year. We also found that flea larvae abundance inside nests was negative and significantly related with brood size. Interactions of fleas with other arthropods may explain this result. In addition, Heeb et al. (2000) did not find a significant effect of brood size on the infestation by fleas in Great tit nests. Additionally, the final abundance of biting midges was positive and significantly related with hatching date at day 13 of nestling age. Previous studies in the same study area have showed that biting midges abundance in blue tit nests increase with hatching date (Tomás et al. 2008b; Martínez de la Puente et al. 2009) probably because conditions for biting midges reproduction improve with season.

Blowfly pupae abundance in nests was negative and significantly related with the condition of blue tit females. Tomás et al. (2005) also found a negative effect of blowfly pupae on the mass of blue tit females in this same area of study. This could be indicative of the assumption of part of the cost of parasitism by female blue tits to reduce effects on nestlings (Tripet and Richner, 1997; Bouslama et al. 2002).

In general, our results suggest that nest boxes provide an isolated environment for both birds and parasites within the complex ecosystem of the forest. Clearly more studies are necessary to better understand the relationships between nest environment and hosts and parasites. In this respect the analysis of gases and their variation inside nests could be of great interest to understand the different relationships between these organisms.

## Acknowledgements

We thank the National Museum of Natural Sciences (MNCN) for providing facilities for this research. Moreover, this study is a contribution to the research developed at the ‘El Ventorrillo’ field station. This study was funded by the project CGL2015-67789-C2-1-P, PGC2018-097426-B-C21 and CGL2016-78318-C2-1-R (MINECO/FEDER). The funders had no role in study design, data collection and analysis, decision to publish, or preparation of the manuscript. The Junta de Castilla y León the authorized the ringing and handling of birds.

